# Cellular Senescence Mediates Doxorubicin Chemotherapy-Induced Aortic Stiffening: Role of Glycation Stress

**DOI:** 10.1101/2025.05.13.653823

**Authors:** Ravinandan Venkatasubramanian, Mary A. Darrah, Sophia A. Mahoney, David A. Hutton, Grace S. Maurer, Katelyn R. Ludwig, Nicholas S. VanDongen, Nathan T. Greenberg, Abigail G. Longtine, Vienna E. Brunt, Parminder Singh, James J. Galligan, Marissa N. Trujillo, Pankaj Kapahi, Simon Melov, Judith Campisi, Matthew J. Rossman, Douglas R. Seals, Zachary S. Clayton

## Abstract

**Background:** Mechanisms underlying Doxorubicin (Doxo) chemotherapy-induced aortic stiffening are incompletely understood.

**Objectives:** Determine the role of cellular senescence and the senescence-associated secretory phenotype (SASP) in mediating Doxo-induced aortic stiffening and the influence of senolytic therapy.

**Methods:** Aortic stiffness (aortic pulse-wave velocity [PWV]), and associated mechanisms were assessed in young adult p16-3MR mice, a model that allows for genetic-based clearance of senescent cells with ganciclovir [GCV]. Young (4-6 month) mice were injected with Doxo and subsequently treated with GCV or the senolytic ABT263. We evaluated the influence of SASP-associated circulating factors in plasma (the circulating SASP milieu) in mediating aortic stiffening *ex vivo* (aortic elastic modulus) and examined the contribution of glycation stress.

**Results:** Doxo increased aortic PWV (425D±D6 vs. control, 353D±D5Dcm/sec; P<0.05), an effect prevented by both GCV (348D±D4Dcm/sec) and ABT263 (342D±D7Dcm/sec; P<0.05 for both vs. Doxo). Plasma from Doxo-treated mice induced aortic stiffening *ex vivo* (P<0.05 vs. plasma from control mice), whereas plasma from Doxo-GCV and Doxo-ABT263 groups did not. Glycation stress was implicated in SASP-mediated aortic stiffening with Doxo, as inhibition of receptor mediated glycation stress signaling attenuated plasma-induced aortic stiffening.

**Conclusion:** Cellular senescence and the circulating SASP milieu contribute to Doxo-induced aortic stiffening. Senolytics hold promise for preserving aortic stiffening following Doxo exposure.

**Translational perspective:** Chemotherapy-induced cardiovascular toxicity is a concern for cancer survivors. This study identifies cellular senescence and the senescence-associated secretory phenotype (SASP) as underlying mechanisms of doxorubicin chemotherapy-induced aortic stiffening – an antecedent to overt cardiovascular disease (CVD). We also provide complementary lines of evidence that glycation stress mediates the mechanistic link between doxorubicin, cellular senescence, the SASP and aortic stiffening. Lastly, we demonstrate the efficacy of senolytic therapy for targeting cellular senescence, the SASP and glycation stress to prevent doxorubicin-induced aortic stiffening. These results offer a novel and clinically actionable approach to preserving vascular health in cancer survivors and mitigating CVD risk.

## INTRODUCTION

Cardiovascular disease (CVD) and cancer remain the top two causes of death globally. In the U.S. alone, over 2 million new cancer cases are projected this year^1^, with nearly 650,000 patients expected to receive chemotherapy treatment^1,2^. While these treatments are often lifesaving, they carry a heavy cost—significant cardiovascular toxicity. Survivors of cancer who have undergone chemotherapy face a markedly higher lifetime risk of developing CVD compared to healthy individuals matched for age and sex^3^. In fact, CVD has emerged as the primary driver of long-term health complications and early death in this population^1,3^. Anthracyclines, first-line chemotherapeutic agents for several common cancers, have toxic effects on the cardiovascular system^4,5^. Doxorubicin (Doxo) is the most widely used anthracycline^6,7^; helps many survive cancer—only for them to face severe cardiovascular complications and premature death from CVD^1,6,^.

The great majority of CVD-related mortality following Doxo treatment stems from clinical atherosclerotic diseases^3^. A major antecedent of these CV disorders is the stiffening of the large elastic arteries (primarily the aorta)^8^. Doxo-associated aortic stiffening, as demonstrated by increased aortic pulse wave velocity (PWV), occurs in part as a result of enhanced pro-inflammatory signaling, increased collagen deposition, elastin degradation, and increased glycation stress, as marked by an increased expression of advanced glycation end products (AGEs)^9^. Glycation stress results from the non-enzymatic modification of proteins and lipids by reactive carbonyl species, leading to the formation of AGEs. These AGEs can stiffen arteries through two primary mechanisms: crosslinking of structural proteins in the vascular wall, and receptor-mediated pathways involving the receptor for AGEs (RAGE)^10^. While the crosslinking effects of AGEs have been more thoroughly studied in the context of vascular stiffening^11^, the direct influence of non-crosslinking AGEs in vascular stiffening — specifically via the AGE-RAGE axis — remains poorly defined^10,12^. Additionally, the mechanistic event(s) governing the role of non-crosslinking AGEs with Doxo-induced aortic stiffening are incompletely understood.

One likely mechanism contributing to Doxo-induced aortic stiffening is cellular senescence^5,13^, a state of cell cycle arrest coupled with the production of pro-inflammatory factors, termed the senescence associated secretory phenotype (SASP)^14^. Physiological levels of cellular senescence are critical for many biological processes (e.g., cancer suppression; wound healing)^15^; however, excessive accumulation of senescent cells occurs in multiple tissues following Doxo treatment^5,13^. This accumulation can induce tissue dysfunction, at least in part through the SASP, and may be involved in select Doxo-induced pathologies. Thus, cellular senescence may contribute to increased aortic stiffness with Doxo. However, the role of cellular senescence and the SASP in mediating Doxo-induced aortic stiffening has not been investigated. Moreover, compounds that selectively clear senescent cells (termed senolytics) improve select indices of physiological dysfunction induced by Doxo chemotherapy^16–18^. Senolytic treatment following Doxo chemotherapy may hold promise for preventing aortic stiffening; however, the impact of senolytics on Doxo-induced aortic stiffening has not yet been assessed.

Here, we performed two complementary studies. In study 1, we utilized the p16-3MR mouse model which allows for genetic-based clearance of excess senescent cells^19^, to establish the role of cellular senescence and the SASP in mediating aortic stiffening with Doxo. In study 2, we targeted cellular senescence with the pharmacological senolytic agent ABT263^18,20^ in mice to establish proof-of-principle efficacy of using senolytic therapy for preventing aortic stiffening with Doxo. We also used a variety of approaches to elucidate the mechanistic role of non-crosslinking-based glycation stress in regulating cellular senescence-mediated aortic stiffening with Doxo

## MATERIAL AND METHODS

### Data Availability

Detailed description of the methods is available under **Supplementary Material**. In the same cohort of mice, we also assessed endothelial function. These data are reported in a companion manuscript ^21^. Select findings from that study are referenced here to provide context for and complement the current analyses.

### Animals

All male and female p16-3MR mice (4–6-month) were housed in a conventional facility on a 12-hour light/dark cycle, given ad libitum access to an irradiated, fixed, and open standard rodent chow (Inotiv/Envigo 7917) and drinking water. *In vivo* testing (blood pressure and aortic PWV) was performed before and 2-4 weeks after the completion of the intervention periods. All mice were euthanized by cardiac exsanguination while maintained under anesthesia (inhaled isoflurane) 2 to 4 weeks following the completion of the intervention periods (allowing for 1 week of recovery and >1 week for *in vivo* post-testing). After cardiac exsanguination, the thoracic aorta was excised, dissected free of perivascular adipose tissue, sectioned, and stored for protein abundance by JESS capillary electrophoresis-based immunoblotting (Protein Simple, San Jose, CA) and immunofluorescence. Investigators were blinded to the treatment group for data collection and biochemical analyses. All animal protocols were approved by the University of Colorado Boulder Institutional Animal Care and Use Committee (protocol no. 2618) and complied with the National Institutes of Health Guide for the Care and Use of Laboratory Animals. Details on all procedures and antibodies are provided in the **Supplemental Material**.

### Study 1: Genetic-based clearance of senescent cells with GCV in p16-3MR mice

p16-3MR mice were bred, weaned, and aged in the animal care facility at the University of Colorado Boulder. At 4 months of age, male and female p16-3MR mice received either a single intraperitoneal (IP) injection of Sham (sterile saline) or Doxo (10 mg/kg in Sham). One week later, mice either received the vehicle (Veh; saline) or ganciclovir (GCV; 25 mg/kg/day in Veh) by IP injection (IP) for 5 consecutive days, which is the standard approach for clearing senescent cells in this model, as we have described previously^20^. This equated to 4 groups/sex: Sham-Veh; Sham-GCV; Doxo-Veh; Doxo-GCV. Throughout this intervention period, 5 mice died as a result of Doxo-related attrition (3 males and 2 females), which resulted in a final sample size of Sham-Veh, n = 22; Sham-GCV, n = 13; Doxo-Veh, n = 23; and DOXO-GCV, n = 25.

### Study 2: Senolytic-based clearance of senescent cells with ABT263 in mice

Male and female p16-3MR mice received a single IP injection of Sham (sterile saline) or Doxo (10 mg/kg in Sham). One week later, mice either received the vehicle (Veh; 10% ethanol, 30% PEG400, 60% Phosal 50 PG) or the senolytic ABT263 (50 mg/kg/day in Veh) by oral gavage on an intermittent one week on; two weeks off, one week on dosing paradigm, as we have previously described^18^. There were 4 groups/sex: Sham-Veh; Sham-ABT263; Doxo-Veh; Doxo-ABT263. Throughout this intervention period, 3 male mice died (two due to Doxo-related attrition and one due to ABT263-related attrition), which resulted in a final sample size of Sham-Veh, n = 11; Sham-ABT263, n = 11; Doxo-Veh, n = 9; and Doxo-ABT263, n = 11.

### Experimental procedures

#### Aortic stiffness and blood pressure

Aortic stiffness was assessed *in vivo* as aortic PWV before and after the dosing period. Aortic PWV in mice is the translational gold standard approach of non-invasively measuring arterial stiffness *in vivo* in humans via carotid-femoral PWV^8,22^. Aortic PWV was conducted via Doppler ultrasonography (Doppler Signal Processing Workstation, Indus Instruments, Webster, TX,) as previously described by our laboratory^18,20,23^. Briefly, mice were placed supine on a heat pad under light inhaled isoflurane anesthesia (1.5%-2.5%) in O_2_. Front and hind limbs were secured to electrodes in order to monitor heart rate, which was maintained between ∼350-500 beats/min. Doppler probes were placed on the transverse aortic arch and abdominal aorta. Three consecutive 2 sec ultrasound tracings were recorded, and time between the R-wave of the ECG to the foot of the Doppler signal (*i*.*e*., average pre-ejection time) was determined for each location. PWV (cm/sec) was calculated by dividing the distance between the two probes by the difference in the pre-ejection times of the aortic arch and abdominal aorta (i.e., Time_abdominal_ – Time_arch_). Blood pressure was measured in vivo over three consecutive days using a noninvasive tail-cuff method (CODA; Kent Scientific, Torrington, CT), as previously described^8,16,18^.

#### Aortic elastic modulus and circulating SASP (plasma)-mediated intrinsic mechanical wall stiffness

Elastic modulus was measured in two 1mm segments of the thoracic aorta as described in the **Supplementary Materials**. For the circulating SASP-induced changes in aortic stiffness, two 1mm segments of thoracic aorta were collected at sacrifice from young adult (4-6 months) intervention-naïve donor mice and cleared of any perivascular adipose and connective tissue. Next, aortic rings from each donor animal were incubated in standard media (DMEM + 1% penicillin-streptomycin) in duplicate for 48 hours in the following conditions: 1) 10% fetal calf serum (control condition); 2) 10% Sham-Veh plasma; 3) 10% Doxo-Veh plasma; 4) 10% Doxo-ABT263 or Doxo-GCV plasma. All plasma was sex-matched to the sex on the donor animal.

After the incubation period, aortic elastic modulus was assessed using the pin myograph (Danish Myo Technology, Denmark) as previously described^20^. Plasma-induced changes in aortic elastic modulus are expressed as a fold-change in comparison to the control incubation condition.

### Statistical Analysis

All details of all statistical analyses are available in the **Supplemental Material**. Unless otherwise noted, data are presented as mean ± SEM in the text, figures, and tables. Statistical significance was defined as α = 0.05. All analyses were conducted using Prism, version 10 (GraphPad Software, Inc., La Jolla, CA).

## RESULTS

### Animal characteristics

#### Study 1: p16-3MR Mouse Study with GCV Administration

To investigate whether cellular senescence directly contributes to Doxo-induced aortic stiffening, we employed the p16-3MR mouse model, a widely accepted tool for studying the impact of p16^INK4A^-related senescence on physiological outcomes^24^. This model enables targeted elimination of p16^INK4A^-expressing senescent cells using the antiviral drug GCV^4,19,20^. In the present study, young male and female p16-3MR mice received either a single IP injection of Sham or Doxo and one week later were either treated with Veh or GCV for 5 consecutive days via IP injection. At the time of sacrifice, animals that received Doxo had significantly lower total body, quadriceps and visceral adipose tissue mass which was not prevented by suppression of excess senescent cells with GCV, as described in a companion manuscript^21^. We observed no group differences in aortic diameter, wall thickness, and blood pressure (**Table 1**). As previously reported in the companion manuscript with this model, Doxo administration results in higher expression of senescence and SASP biomarkers in the aorta, which were prevented by GCV treatment—confirming effective suppression of senescent cells in the aorta with this study paradigm^21^.

**Table 1.** Characteristics of mice in the doxorubicin (DOXO) GCV study (**Study 1**)

|  | **Sham**<br>**Veh** | **Sham**<br>**GCV** | **DOXO**<br>**Veh** | **DOXO**<br>**GCV** |
| --- | --- | --- | --- | --- |
| ***n*** | 27<br>*10 female*<br>*17 male* | 20<br>*9 female*<br>*11 male* | 18<br>*9 female*<br>*9 male* | 26<br>*14 female*<br>*12 male* |
| **Aorta** |  |  |  |  |
| **Diameter, $\mu\text{m}$** | 601 ± 15 | 624 ± 25 | 611 ± 17 | 627 ± 28 |
| **Intima media thickness, $\mu\text{m}$** | 41 ± 1 | 39 ± 3 | 43 ± 2 | 43 ± 4 |
| **Systolic blood pressure, mmHg** |  |  |  |  |
| **Pre** | 104 ± 2 | 106 ± 1 | 103 ± 2 | 105 ± 1 |
| **Post** | 103 ± 2 | 106 ± 2 | 105 ± 3 | 103 ± 2 |
| **Diastolic blood pressure, mmHg** |  |  |  |  |
| **Pre** | 75 ± 1 | 78 ± 2 | 74 ± 2 | 76 ± 1 |
| **Post** | 75 ± 2 | 78 ± 1 | 76 ± 2 | 73 ± 2 |

Data are mean ± SEM.

#### Study 2: Senolytic (ABT263) Administration Mouse Study

To test the pharmacological effects of senolytics on targeting excess cellular senescence to prevent Doxo-induced aortic stiffening, we used the well-established senolytic drug ABT263, a well-characterized synthetic senolytic that we recently showed can reduce senescent cell burden and lower aortic stiffness in aged mice^20^, and applied the same approach in the present study. ABT263 works by inhibiting Bcl-2, an anti-apoptotic protein upregulated in senescent cells, thereby selectively triggering apoptosis in senescent cells^17^. Here, we used young male and female p16-3MR mice. One week after a single IP injection of Sham or Doxo we treated animals with either Veh or ABT263. At the time of sacrifice, animals that were administered with Doxo had lower visceral adipose tissue mass relative to Sham-administered animals, as shown in the companion manuscript^21^. We observed no group differences in aortic diameter and wall thickness, or blood pressure (**Supplemental Table 1**). Consistent with our prior findings, Doxo increased biomarkers of cellular senescence and the SASP in the aorta, all of which were prevented by ABT263 treatment. Additionally, in this model, Doxo increased aortic Bax:Bcl2 ratio—an established surrogate marker of pro-apoptotic signaling in senescent cells^17^—which was prevented by ABT263 treatment^21^, confirming molecular target engagement of ABT263. The model verification data for both the studies is reported in the companion manuscript^21^.

#### Aortic stiffness

We have previously shown that Doxo administration can cause aortic stiffening^9^. Thus, here we sought to determine whether excessive cellular senescence contributes to aortic stiffening with Doxo. To accomplish this goal, we first assessed aortic PWV in study 1 prior to Doxo/Sham administration and after the cessation of treatment in vehicle-and GCV-treated animals. Aortic PWV was 17% higher following Doxo administration (Doxo-Veh, pre [362 ± 5 cm/sec] vs. post [425 ± 6 cm/sec] treatment, *P* < 0.0001), which was prevented with GCV treatment (Doxo-GCV, pre [350 ± 4 cm/sec] vs. post [348 ± 4 cm/sec] treatment, *P* = 0.99; Doxo-Veh post-treatment vs. Doxo-GCV post-treatment, *P* < 0.0001; **Figure 1A**). These effects occurred independent of any change in systolic or diastolic blood pressure (**Table 1**).

**Figure 1.**
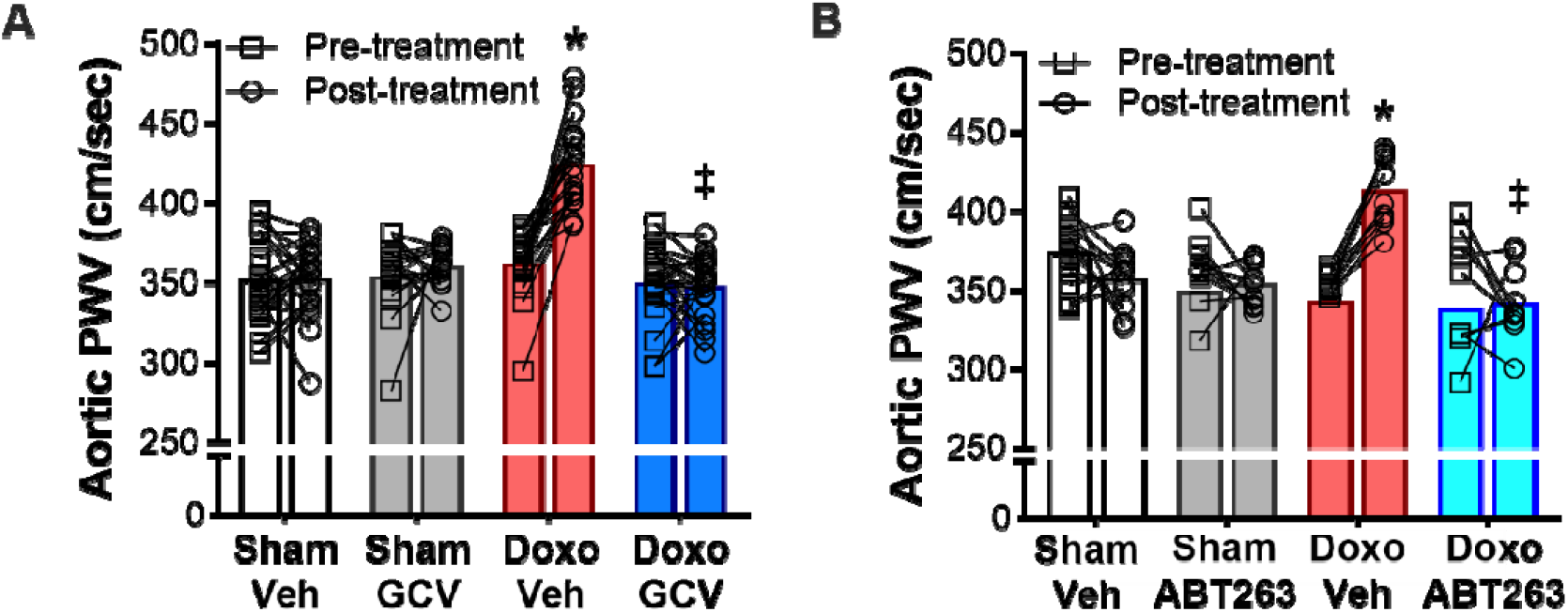
Doxorubicin (Doxo)-induced cellular senescence mediates aortic stiffening which is prevented by senolytic treatment. Aortic pulse wave velocity (PWV) pre-and post-treatment in study 1 **(A)** and 2 **(B)**. All values are mean. n=13-25 in study 1 PWV, n=9-12 in study 2 PWV. *p<0.05 vs Sham-Veh, ‡p<0.05 vs Doxo-Veh.

Next, in study 2 we sought to determine the efficacy of senolytic treatment in preventing Doxo-induced aortic stiffening by assessing aortic PWV prior to Doxo/Sham administration and one week after the cessation of treatment in vehicle-and ABT263-treated animals. Similar to study 1, aortic PWV was 17% higher following Doxo administration (Doxo-Veh, pre [343 ± 11 cm/sec] vs. post [404 ± 15 cm/sec] treatment, *P* < 0.0001), which was prevented with ABT263 treatment (Doxo-ABT263, pre [339 ± 16 cm/sec] vs. post [341 ± 7 cm/sec] treatment, *P* = 0.99; Doxo-Veh post-treatment vs. Doxo-GCV post-treatment, *P* < 0.0001; **Figure 1B**). These effects occurred independent of changes in systolic or diastolic blood pressure (**Supplementary Table 1**). Taken together these results indicate that cellular senescence underlies the increase in aortic stiffness with Doxo and senolytic therapy can prevent Doxo-induced aortic stiffening independent of any change in systolic or diastolic blood pressure.

### Aortic wall remodeling

To confirm our PWV results we then sought to determine if there were any adverse changes in the abundance of the arterial wall structural proteins type-1 collagen and elastin fragmentation. Type-1 collagen is a component of the arterial adventitia that provides rigidity/stiffness to the arterial wall, whereas *α*-elastin is a protein that confers arterial wall elasticity^14,25^. The aortic abundance of type-1 collagen and elastin fragmentation were only assessed in study 2 given that difference in aortic intrinsic wall stiffness was similar in studies 1 and 2. We found that there were no differences between groups in the abundance of type-1 collagen (**Supplementary Figure 1A**) or elastin fragmentation (as measured by the number of elastin breaks) (**Supplementary Figure 1B**). Together, these results suggest that cellular senescence induces aortic stiffening with Doxo via mechanisms that are independent of changes in arterial wall structural proteins indicating that other factors may contribute.

### Cellular senescence mediated changes in the circulating SASP milieu influence vascular function: prevention by senolytic therapy

To better capture the *in vivo* environment of the aorta, we next sought to determine the role of humoral factors in plasma as mediators of aortic stiffening with Doxo, given that SASP factors are secreted in the circulation (e.g., into the plasma)^25,26^ can exert continuous and direct effects on the arterial wall^20^. Specifically, we recently demonstrated that circulating SASP factors in plasma, in part, mediate aortic stiffening with chronological aging^20^. To accomplish this, we exposed aortic rings, isolated from young adult (6 months) intervention naïve p16-3MR female and male mice to standard cell culture media containing either fetal calf serum (control) or serum-free media supplemented with 10% sex-matched plasma from groups in study 1 (Sham-Veh, Doxo-Veh and Doxo-GCV), and performed the aortic elastic modulus assay, as described above. We found that plasma from Doxo-Veh mice evoked a 2-fold increase in aortic elastic modulus (*P* = 0.02 vs. control media), whereas there was no increase in elastic modulus with plasma from Doxo-GCV mice (Doxo-Veh vs. Doxo-GCV mice, *P* = 0.003; Doxo-GCV vs. control media, *P* = 0.72 **Figure 2A**). Moreover, there was no difference between Sham-Veh plasma and control media (*P* = 0.27; **Figure 2A**). Together, these results suggest that SASP factors in plasma likely contribute to the increase in aortic stiffness following Doxo administration. In further support of this concept, linear regression analyses showed that plasma-mediated changes in aortic elastic modulus were related to post-intervention aortic PWV values *in vivo* (R^2^ = 0.28, *P* = 0.028; **Figure 2C**).

**Figure 2.**
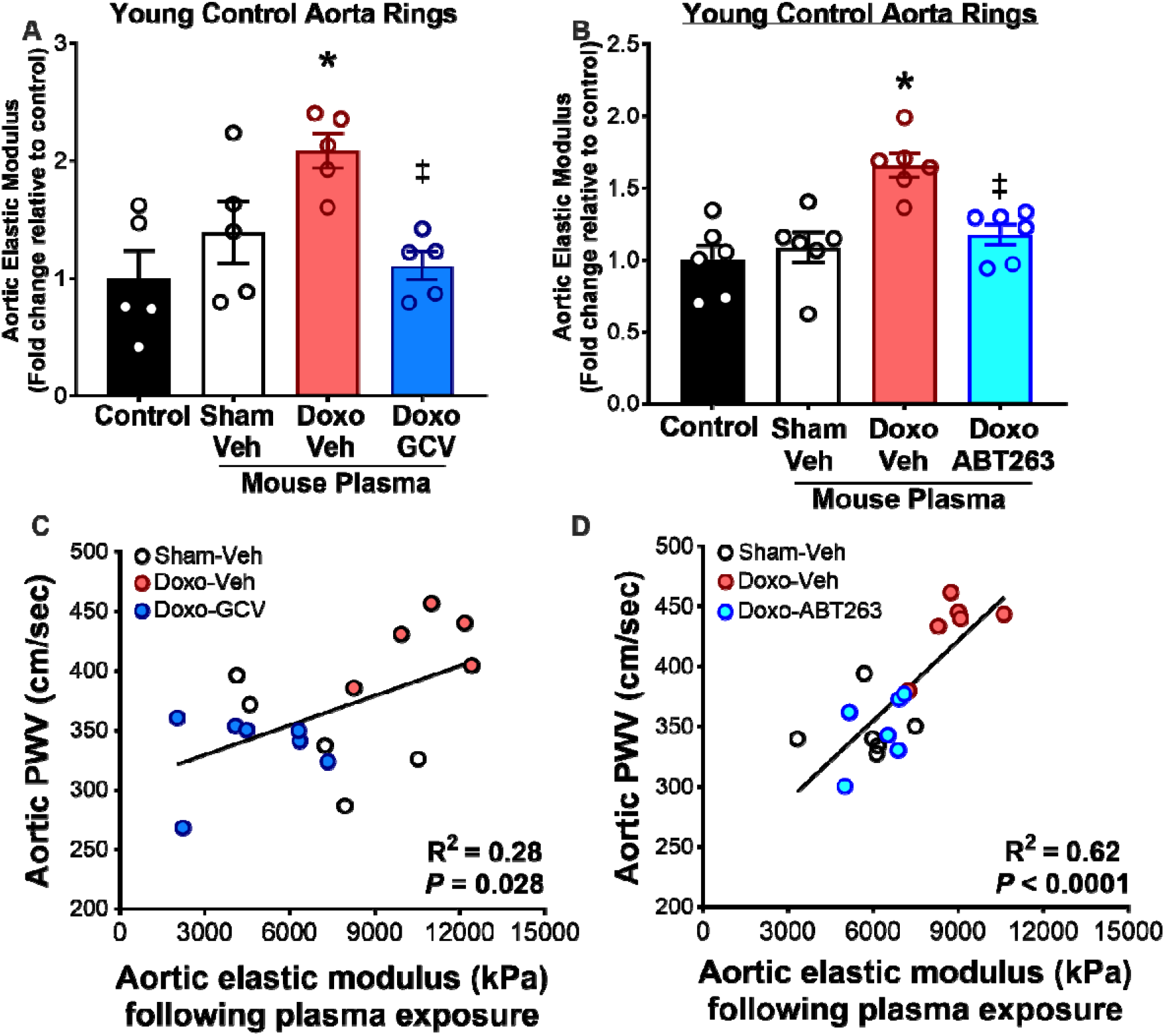
Doxo-mediated cellular senescence alters circulating senescence associated secretory phenotype (SASP) milieu causing vascular dysfunction which is prevented by senolytic treatment. Changes in elastic modulus of aorta rings from young intervention naïve mice when exposed to plasma from mice in study 1 **(A)** and 2 **(B)**. Aortic elastic modulus following plasma exposure relative to the post-intervention aortic pulse wave velocity (PWV) value of plasma donor mouse in study 1 **(C)** and 2 **(D)**. Values in **(A)** and **(B)** are mean ± SEM. n=5/group in study 1. n=6/group in study 2. *p<0.05 vs Sham-Veh, ‡p<0.05 vs Doxo-Veh.

Following, we performed a similar set of experiments using plasma samples from study 2, to determine if targeting excess cellular senescence with senolytic therapy could mitigate aortic stiffening induced by the circulating SASP milieu following Doxo administration. Similar to study 1, we found that plasma from Doxo-Veh mice evoked an ∼2-fold increase in aortic elastic modulus (*P* = 0.013 vs. control media), whereas there was no increase in elastic modulus with plasma from Doxo-ABT263 mice (Doxo-Veh vs. Doxo-ABT263 mice, *P* = 0.009; Doxo-ABT263 vs. control media, *P* = 0.26; **Figure 2B**,). Moreover, there was no difference between Sham-Veh plasma and control media (*P* = 0.56; **Figure 2B**). We also found that plasma-mediated changes in aortic elastic modulus were related to post-intervention aortic PWV values *in vivo* (R^2^ = 0.62, *P* < 0.0001; **Figure 2D**). Collectively, these results suggest that the circulating SASP milieu may directly influence aortic stiffening following Doxo administration, and that prevention of unfavorable changes to the circulating SASP milieu may be a mechanism by which senolytic therapy prevents Doxo-induced aortic stiffening.

### Cellular senescence underlies the Doxo-induced increase in aortic advanced glycation end products (AGEs): Influence of the circulating SASP milieu

#### Circulating Methylglyoxal derived hydroimidazalone-1 (MGH-1)

Another component of the aortic wall that can modulate stiffening is AGEs^12,28^. AGEs are formed via irreversible reactions produced through the non-enzymatic glycation of proteins, lipids, and nucleic acids, playing a pivotal role in mediating arterial stiffness^8,14,29^ via structural and functional mechanisms. AGEs can structurally alter stiffness by cross-linking extracellular matrix proteins (non-receptor mediated) or they can functionally alter stiffness by lowering nitric oxide bioavailability, increasing inflammation and oxidative stress. This occurs through AGEs binding to the receptor for AGE (RAGE)^10,12,28^. Among the various AGEs, methylglyoxal-derived hydroimidazolone-1 (MGH-1) is a key biomarker of glycation stress and is implicated in aortic stiffening and CVD^29–31^. While cross-linking AGEs (non-receptor mediated) have been well-studied for their role in directly inducing aortic stiffening through structural modifications of the extracellular matrix^11^, there is limited direct evidence linking non-crosslinking AGEs, such as MGH-1, to aortic stiffening. The present study aimed to address this critical gap in knowledge by investigating how non-crosslinking AGEs like MGH-1 contribute to aortic stiffening via activation of the AGE-RAGE signaling pathway, offering new insights into the mechanisms of aortic stiffening that may differ from those induced by cross-linking AGEs.

Via plasma proteomic analyses, we recently demonstrated that glycolytic metabolism was the pathway most influenced with aging and clearance of excess senescent cells^20^. We reasoned that in states of mitochondrial dysfunction (*due to* excess cellular senescence), that there is a greater reliance on glycolysis^32^. Additionally, we have also previously shown that Doxo administration increases aortic abundance of AGEs^9^. Thus, given our observations of mitochondrial dysfunction in our previous series of studies (*i*.*e*., excessive mtROS production in arteries following systemic Doxo administration)^21,23^, we first leveraged our plasma exposure model and sought to determine if exposure of aorta rings from intervention naïve p16-3MR mice to sex-matched plasma from animals in study 1 led to differential expression of MGH-1. Indeed, we found that exposure of plasma from Doxo-Veh mice led to an increase in aortic MGH-1 (*P* = 0.023 vs. Sham-Veh), which was prevented with plasma from Doxo-GCV animals (*P* = 0.47 vs. Sham-Veh; **Figure 3A**). We then sought to determine if plasma from animals in study 2 could induce similar changes in aortic MGH-1, as observed with plasma from study 1. Exposure of naïve aorta rings to plasma from Doxo-Veh mice led to an increase in aortic MGH-1 abundance (*P* = 0.008 vs. Sham-Veh), which was fully prevented with plasma from Doxo-ABT263 animals (*P* = 0.45 vs. Sham-Veh; Doxo-Veh vs. Doxo-ABT263, *P* = 0.004; **Figure 3B**). Following, we interrogated the relation between plasma-induced aortic MGH-1 and plasma-induced aortic elastic modulus *ex vivo* by performing linear regression analyses. These analyses demonstrated that aortic MGH-1 was positively associated with plasma-induced aortic elastic modulus *ex vivo* (R^2^ = 0.17, *P* = 0.12 in study 1 and R^2^ = 0.21, *P* = 0.054 in study 2; **Supplementary Figure 2**). Together, these findings suggest that circulating factors induced by Doxo promote aortic stiffening, at least in part, by increasing aortic MGH-1—linking SASP signaling to AGE accumulation and vascular dysfunction.

**Figure 3:**
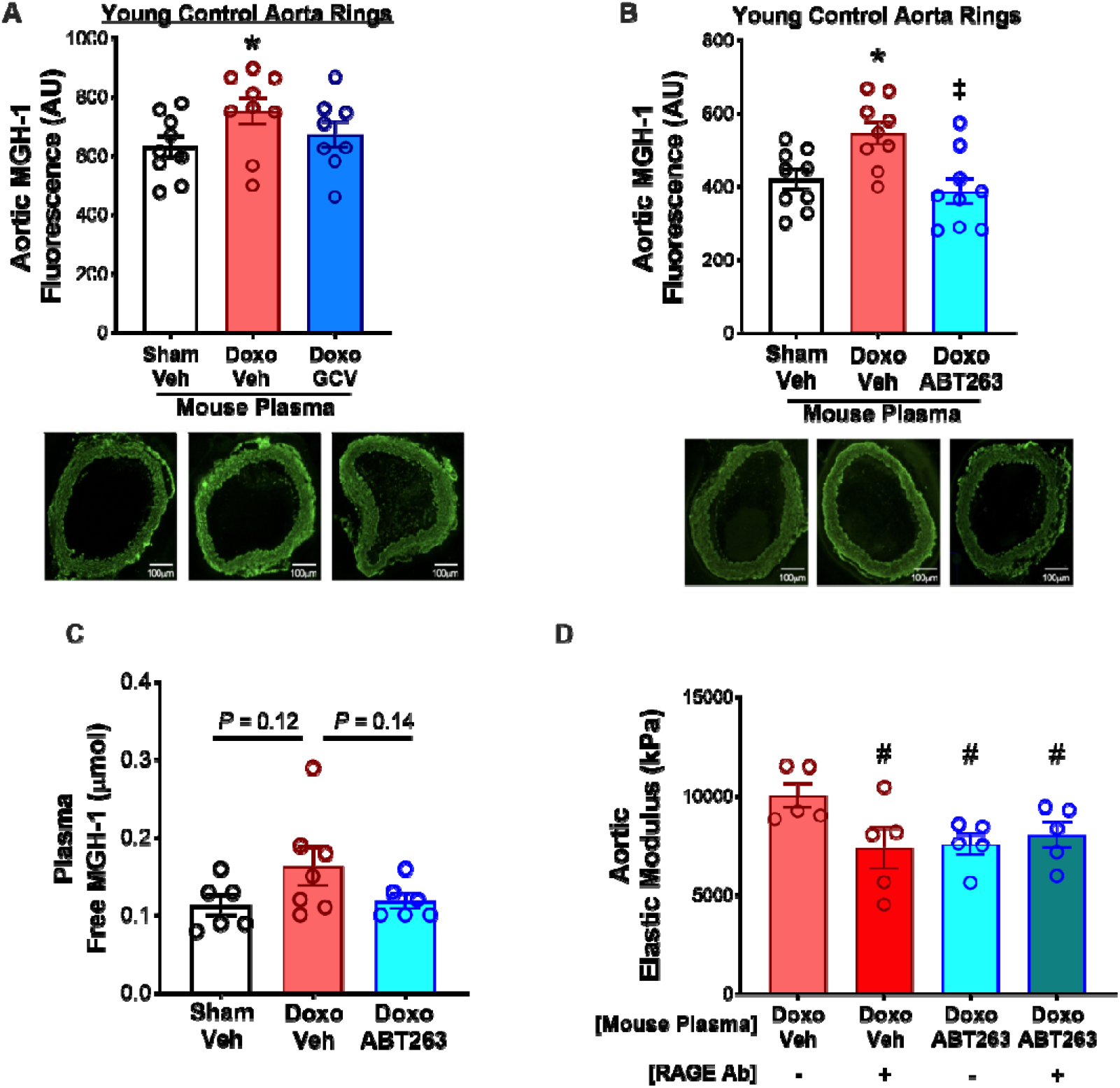
Doxo-induced circulating SASP milieu increase aortic methylglyoxal derived hydroimidazolone-1 (MGH-1) accumulation. Immunofluorescence (IF) staining for MGH-1 in intervention naïve aorta rings exposed to plasma from mice that received treatment **(A)**. IF staining for MGH-1 in intervention naïve aorta rings exposed to plasma collected from mice that received treatment **(B)**. Circulating MGH-1 levels from mice in study 2 **(C)**. Changes in elastic modulus of aorta rings from young intervention naïve mice when exposed to plasma from mice in study 2 in the presence/absence of RAGE antibody (Ab) **(D)**. All values are mean ± SEM. n=9/group/study for plasma-induced MGH-1 IF. n=6-7/group for plasma free MGH-1 levels. n=5/group for aortic intrinsic mechanical wall stiffness with RAGE Ab. *p<0.05 vs Sham-Veh, ‡p<0.05 vs Doxo-Veh, #p<0.05 vs Doxo-Veh in absence of RAGE Ab. AU: arbitrary unit.

Given the observed increases in aortic MGH-1 following plasma exposure, we next measured circulating levels of MGH-1 in study 2. For comparison, we also assessed select additional AGEs implicated in CVD^30^—including carboxy-ethyl-lysine (CEL), carboxy-ethyl-arginine (CEA), and carboxy-methyl-lysine (CML) (**Supplementary Table 2**)—none of which were significantly altered. Among the AGEs studied, we found that only MGH-1 was altered with Doxo administration and senolytic therapy. Doxo-Veh mice tended to have a higher circulating level of MGH-1 (*P* = 0.12 vs. Sham-Veh) which was not present in plasma from Doxo-ABT263 animals (*P* = 0.76 vs. Sham-Veh; **Figure 3C**). Moreover, plasma MGH-1 tended to be lower in Doxo-ABT263 mice relative to Doxo-Veh mice (*P* = 0.14; **Figure 3C**).

Given that circulating MGH-1 was the only AGE that was generally different between groups in study 2, and it primarily exerts its effects via the RAGE-AGE signaling pathway^12^, we next sought to determine the cause-and-effect role of circulating AGEs in mediating aortic stiffening with Doxo administration. Plasma exposure experiments revealed clear evidence of circulating SASP milieu-mediated aortic stiffening. When aorta rings of intervention naïve young adult p16-3MR mice were incubated with sex-matched plasma from animals in study 2, we again observed lower aortic elastic modulus following exposure to plasma from Doxo-ABT263 mice relative to Doxo-Veh animals (P = 0.041; **Figure 3D**). Blocking AGE-RAGE signaling with a RAGE antibody significantly reduced the elastic modulus in aorta rings exposed to Doxo-Veh plasma, bringing it down to levels observed with Doxo-ABT263 plasma. In contrast, RAGE inhibition had no further effect in the Doxo-ABT263 plasma group, suggesting that AGE-RAGE signaling mediates the differential stiffening response (Doxo-Veh + RAGE Ab vs Doxo-ABT263, P=0.911, **Figure 3D**). These data indicate that the circulating SASP milieu following Doxo administration promotes aortic stiffening through RAGE signaling, and that inhibition of excessive RAGE activation is a likely mechanism by which senolytic therapy protects against aortic stiffening. It also suggests that factors in the circulating SASP milieu, such as MGH-1, contribute to aortic stiffening by promoting non-structural changes in the aorta.

### Senolytic therapy following Doxo administration prevents elevated Rage and MGH-1 expression independent of Glyoxalase-1 (Glo-1)

Given our findings with influence of AGE-RAGE signaling in mediating aortic stiffening with Doxo administration and prevention with senolytic therapy, we next sought to determine mRNA expression of *Rage* in aortas from animals in study 2, as *Rage* gene expression is predictive of RAGE protein content^33–35^ We found that Doxo-Veh animals had 2-fold higher *Rage* expression relative to Sham-Veh animals (*P* = 0.045), which was fully prevented with senolytic therapy (Doxo-ABT263 vs. Doxo-Veh, *P* = 0.01; Doxo-ABT263 vs. Sham-Veh, *P* = 0.87; **Figure 4A**).

**Figure 4.**
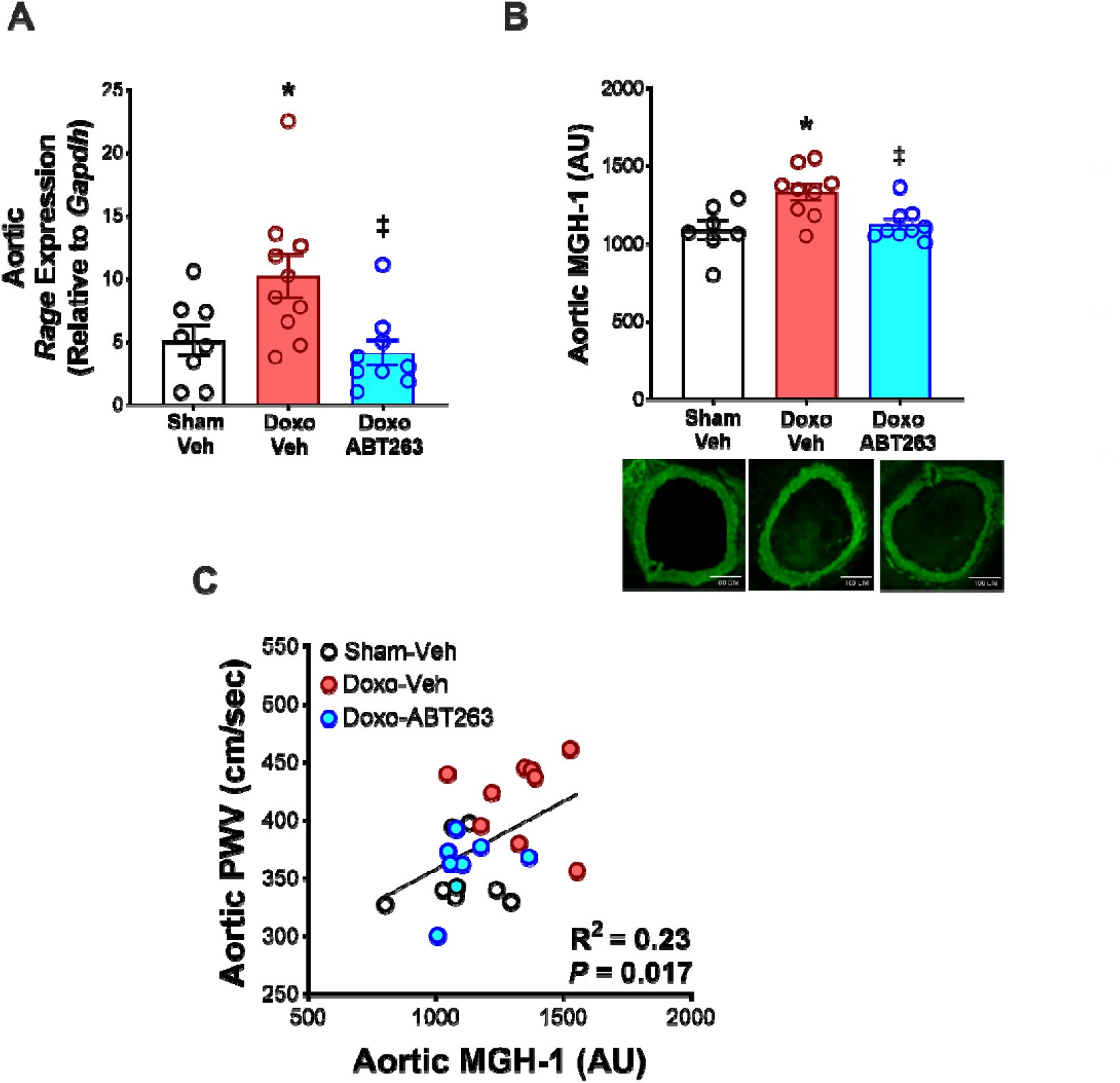
Senolytic therapy following Doxo administration prevents elevated *Rage and MGH-1* expression. Aortic mRNA expression of *Rage* from mice in study 2 **(A)**. Aortic MGH-1 levels from mice in study 2 **(B)**. Aortic MGH-1 levels relative to the post-intervention aortic pulse wave velocity (PWV) value of mice in study 2 **(C)**. All values are mean ± SEM. n=8-10/group for *Rage* expression. n=7-9/group in study 2 for aortic MGH-1 levels. *p<0.05 vs Sham-Veh, ‡p<0.05 vs Doxo-Veh, AU: arbitrary unit

Based on these findings we next aimed to determine if aortic abundance of MGH-1 was elevated following Doxo administration and influenced by senolytic therapy. To achieve this goal, we assessed aortic MGH-1 at time of sacrifice, from animals in study 2. Doxo administration resulted in ∼20% higher aortic MGH-1 abundance (Doxo-Veh vs. Sham-Veh, *P* = 0.009) which was fully prevented with senolytic therapy (Doxo-ABT263 vs. Sham-Veh, *P* = 0.89; Doxo-ABT263 vs. Doxo-Veh, *P* = 0.01) (**Figure 4B**). The increased abundance of MGH-1 remains relevant, as our previous plasma exposure model demonstrated that non-crosslinking AGEs, such as MGH-1, can independently promote aortic stiffening through AGE-RAGE signaling. Following, we interrogated the relation between aortic MGH-1 and post-intervention *in vivo* aortic stiffness by performing linear regression analyses. These analyses demonstrated that aortic MGH-1 abundance was positively and significantly associated with post-intervention aortic PWV (R^2^ = 0.23, *P* = 0.017; **Figure 4C**). These findings confirm that non-crosslinking AGEs, such as MGH-1, can directly mediate aortic stiffening through AGE-RAGE signaling. The positive correlation between MGH-1 abundance and post-intervention PWV further supports this mechanism, providing evidence for the role of non-crosslinking AGEs in aortic stiffening following Doxo administration.

Glyoxalase-1 (Glo-1) is an essential enzyme of the glyoxalase system that plays a critical role in detoxifying methylglyoxal (MGO) (a major precursor for MGH-1), thereby limiting AGE formation and its associated vascular complications^29,36,37^. Lastly, we examined the protein expression of Glo-1 from mice in study 2, and found that Doxo administration significantly lowered aortic Glo-1 expression (Sham-Veh vs Doxo-Veh; *P* = 0.043) which was not influenced by senolytic treatment (Sham-Veh vs Doxo-ABT263, P = 0.047; **Supplementary Figure 3**) suggesting that increased aortic abundance of MGH-1 likely contributes to aortic stiffening with Doxo, while senolytic therapy may mitigate this effect by preventing Doxo-induced increases in MGH-1 levels, rather than via preservation of Glo-1-mediated MGH-1 detoxification.

## DISCUSSION

CVD is a leading cause of death in Doxo chemotherapy-treated cancer survivors^1^. In this study, we used complementary *in vivo* and *ex vivo* experimental models to show that clearing senescent cells after Doxo administration prevents Doxo-induced aortic stiffening. Using the p16-3MR mouse model, which enables genetic clearance of senescent cells with GCV, we first identified a causal role for excess cellular senescence in mediating Doxo-induced aortic stiffening. We then demonstrated that pharmacological clearance of senescent cells with the senolytic agent ABT263 prevented aortic stiffening, highlighting cellular senescence as a target mechanism for preserving aortic stiffness following administration of Doxo chemotherapy.

Stiffening of the aorta, as indicated by increased PWV, is an independent risk factor for hypertension^8^, kidney disease^38^, cognitive impairment^39^ and risk for diabetes^38^. Human cancer survivors previously treated with Doxo chemotherapy exhibit higher aortic stiffness compared to age-and sex-matched healthy controls^3^; however, the underlying mechanisms mediating aortic stiffening with Doxo are largely unknown, highlighting the importance for interrogating these potential processes. Cellular and molecular mechanisms contributing to increased intrinsic aortic wall stiffness include: (a) greater deposition of collagen-1, the main load-bearing protein in the aortic wall; (b) increased elastin fragmentation, which reduces aortic elasticity; and (c) accumulation of AGEs, which promote aortic stiffening via activation of receptor dependent and receptor independent pathways^30,35^. Additionally, the composition of humoral factors, specifically the SASP, can influence arterial wall stiffening as factors in circulation come in direct and frequent contact with the arterial wall.

In the present study, we show that Doxo-induced cellular senescence mediates aortic stiffening and that senolytic treatment prevents aortic stiffening with Doxo independent of collagen deposition and elastin fragmentation. Moreover, we demonstrate that cellular senescence and the circulating SASP milieu (humoral factors in plasma) from Doxo-treated mice directly promotes aortic stiffening, as this phenotype was prevented when we assessed the effect of plasma from Doxo-treated mice that had been genetically cleared of senescent cells and from mice that had been treated with a senolytic.

Increased glycation stress, as indicated by higher AGE accumulation, is linked to premature and accelerated CV aging and heightened CVD risk^29,30^. Previous studies have demonstrated evidence of increased systemic glycation stress in cancer survivors previously treated with Doxo chemotherapy^41^. Here, we show that Doxo administration increases accumulation of the AGE; MGH-1, which was prevented with senolytic treatment. We also found that MGH-1 is upregulated in the circulation of Doxo-treated mice. Plasma from Doxo-treated mice, but not plasma from Doxo-treated mice cleared of senescent cells, induced MGH-1 accumulation in the aorta of young, intervention-naïve mice, suggesting that cellular senescence with Doxo mediates the aortic stiffening effects via increase in glycation stress. Lastly, we mechanistically show that circulating SASP factors induced by Doxo exert effects on aortic stiffening via the AGE-RAGE pathway.

These findings provide novel insights into the interaction between glycation stress and Doxo-induced cellular senescence, elucidating how these events mediate aortic stiffening. Notably, multiple clinical trials aimed at breaking AGE crosslinks, via the administration of crosslink breaker ALT-711, have yielded inconclusive or modest effects on vascular stiffness^42–44^, raising questions about whether crosslinking is the primary driver. Our data provide insight into these critical gaps in knowledge by demonstrating that a non-crosslinking AGE, MGH-1, contributes to aortic stiffening through AGE-RAGE signaling—a distinct, pro-inflammatory pathway upregulated in response to Doxo administration and prevented by senolytic therapy. This mechanistic link positions non-crosslinking AGE-RAGE signaling, rather than crosslink accumulation alone, as a viable therapeutic target in preventing aortic stiffening in Doxo chemotherapy-treated cancer survivors.

Currently, several clinical trials are underway to evaluate the potential of senolytic therapies to improve physiological function in cancer survivors. For example, one trial (NCT04733534) is testing whether reducing senescent cell burden with the senolytic cocktail dasatinib and quercetin, or with the flavonoid senolytic fisetin, can mitigate frailty in adult survivors of childhood cancer. Another trial (NCT05595499) is investigating the impact of fisetin on physical function in survivors of stage I–III breast cancer. While these studies represent important steps toward clinical translation of senolytic therapy, there are no ongoing trials specifically examining the impact of senolytics on large artery stiffness in cancer survivors. Considering aortic stiffness plays a pivotal role in the development of CVD and hypertension following cancer treatment, future research is needed to determine whether senolytic therapies can mitigate large artery stiffening in Doxo chemotherapy-treated cancer survivors.

In conclusion, this study provides key proof-of-principle evidence that cellular senescence and the circulating SASP milieu play critical mechanistic roles in Doxo-induced aortic stiffening and that senolytic treatment following Doxo administration can prevent these adverse changes to the aorta. Importantly, we demonstrate the role of cellular senescence in mediating additional mechanisms underlying Doxo-induced aortic stiffening, including glycation stress and activation of the AGE-RAGE pathway. Notably, lack of changes in the structural components of the aortic wall (i.e. collagen deposition and elastin fragmentation) might suggest the role of adverse signaling mechanisms in mediating aortic stiffening with Doxo rather than overt structural changes.

