## Supplementary material for "Cellular Senescence Mediates Doxorubicin Chemotherapy-Induced Aortic Stiffening: Role of Glycation Stress": Supplemetary materials

**SUPPLEMENTAL MATERIALS**

**Animals**

All male and female mice were housed in a conventional facility on a 12-hour light/dark cycle, given ad libitum access to an irradiated, fixed, and open standard rodent chow (Inotiv/Envigo 7917) and drinking water. *In vivo* testing (blood pressure and aortic PWV) was performed before and 2-4 weeks after the completion of the intervention periods. All mice were euthanized by cardiac exsanguination while maintained under anesthesia (inhaled isoflurane) 2 to 4 weeks following the completion of the intervention periods (allowing for 1 week of recovery and 1 week for *in vivo* post-testing). After cardiac exsanguination, the thoracic aorta was excised, dissected free of surrounding perivascular adipose and connective tissue, sectioned, and stored in for later protein abundance by JESS capillary electrophoresis-based immunoblotting (Protein Simple, San Jose, CA) and immunofluorescence. Investigators were blinded to the treatment group for data collection and biochemical analyses. All animal protocols were approved by the University of Colorado Boulder Institutional Animal Care and Use Committee (protocol no. 2618) and complied with the National Institutes of Health Guide for the Care and Use of Laboratory Animals.

**Study 1: Genetic-based clearance of senescent cells with GCV in p16-3MR mice**

p16-3MR mice were bred, weaned, and aged (to 4 months of age) in the animal care facility at the University of Colorado Boulder. At 4 months of age, male and female p16-3MR mice received either a single intraperitoneal injection of Sham (sterile saline) or Doxo (R&D Systems #2252/50 Minneapolis, MN) (10 mg/kg in Sham). One week later, mice either received the vehicle (Veh; saline) or ganciclovir (Sigma-Aldrich #G2536, St. Louis, MO) (GCV; 25 mg/kg/day in Veh) by intraperitoneal injection (IP) for 5 consecutive days, which is the standard approach for clearing senescent cells in this model, as we have described previously^1^. This equated to 4 groups/sex: Sham-Veh; Sham-GCV; Doxo-Veh; Doxo-GCV. Throughout this intervention period, 5 mice died as a result of Doxo-related attrition (3 males and 2 females), which resulted in a final sample size of Sham-Veh, n = 22; Sham-GCV, n = 13; DOXO-Veh, n = 23; and DOXO-GCV, n =25 .

**Study 2: Senolytic-based clearance of senescent cells with ABT263 in mice**

Male and female p16-3MR mice received a single intraperitoneal injection of Sham (sterile saline) or Doxo (10 mg/kg in Sham). One week later, mice either received the vehicle (Veh; 10% EtOH, 30% PEG400, 60% Phosal 50 PG) or the senolytic ABT263 (Selleckchem #S1001, Houston, TX) (50 mg/kg/day in Veh) by oral gavage on an intermittent one week on; two weeks off, one week on dosing paradigm, as we have previously described^1,2^. There were 4 groups/sex: Sham-Veh; Sham-ABT263; Doxo-Veh; Doxo-ABT263. Throughout this intervention period, 5 mice died because of Doxo-related attrition, which resulted in a final sample size of Sham-Veh, n = 11; Sham-ABT263, n = 11; Doxo-Veh, n = 9; and Doxo-ABT263, n = 11.

**Experimental procedures**

***Aortic stiffness and blood pressure****.* Aortic stiffness was assessed *in vivo* as aPWV before and after the dosing period. aPWV is translational to the gold standard approach of measuring arterial stiffness in humans, which is via carotid-femoral PWV^5^. aPWV was conducted via Doppler ultrasonography (Doppler Signal Processing Workstation, Indus Instruments, Webster, TX,) as previously described by our laboratory^2,6^. Briefly, mice were placed supine on a heat pad under light inhaled isoflurane anesthesia (1.5%-2.5%) in O_2_. Front and hind limbs were secured to electrodes in order to monitor heart rate, which was maintained between ~350-500 beats/min. Doppler probes were placed on the transverse aortic arch and abdominal aorta. Three consecutive 2 sec ultrasound tracings were recorded, and time between the R-wave of the ECG to the foot of the Doppler signal (*i.e.,* average pre-ejection time) was determined for each location. PWV (cm/sec) was calculated by dividing the distance between the two probes by the difference in the pre-ejection times of the aortic arch and abdominal aorta (i.e., Time_abdominal_ – Time_arch_). To examine the potential contribution of changes in arterial blood pressure to any treatment-related differences in aortic PWV, systolic and diastolic blood pressures were assessed using a CODA noninvasive tail-cuff system (Kent Scientific, Torrington, CT), as we have described previously^2,4^. Briefly, the pressure measurements from 20 collection cycles (following 5 acclimation cycles) over 3 consecutive days were averaged for each mouse at each timepoint.

***Sacrifice and tissue collection****.* Mice were sacrificed using a method approved under the American Veterinary Medical Association guidelines. Mice were anesthetized under inhaled anesthesia (open-drop method) and euthanized via cardiac exsanguination. The heart was removed, cleaned, and weighed. The aorta was excised and rinsed in physiological saline solution (PSS), cleared of perivascular adipose tissue, and sectioned and stored as described below.

***Plasma-mediated intrinsic mechanical wall stiffness****.* For the plasma-induced changes in aortic stiffness, two 1mm segments of thoracic aorta were collected at sacrifice from intervention-naïve mice and cleared of any perivascular connective tissue. The aortic rings were incubated in standard media (DMEM + 1% penicillin-streptomycin) in duplicate for 48 hours in the following conditions: 1) 10% fetal calf serum (control condition); 2) 10% Sham-Veh plasma; 3) 10% Doxo-Veh plasma; 4) 10% Doxo-ABT263/GCV plasma. After the incubation period, elastic modulus was assessed using the pin myograph (Danish Myo Technology, Denmark) as previously described^6^. Plasma-induced changes in aortic elastic modulus are expressed as a fold-change in comparison to the control incubation condition. For the RAGE blocking assay, intervention naiive mouse aortas were incubated with plasma either Doxo-Veh or Doxo-ABT263 mice in presence/absence of 10ug/ml of RAGE antibody (R&D systems #AF1145 Minneapolis, MN). RAGE antibody concentrations were based on previous research^7,8^.

In short, aorta samples were placed in heated (37°C) baths filled with calcium-free, phosphate-buffered saline (PBS). The samples were then mounted on two wire prongs (Danish Myo Technology, Denmark) and pre-stretched for three minutes to a 1mm luminal diameter displacement that was returned to the non-stretched baseline, and this was repeated two more times. Once pre-stretching was complete, aortic ring diameter was increased until 1mN of force was reached and incrementally increased by 50 mN every 3 minutes thereafter until failure (i.e., the ring broke). The force corresponding to each stretching interval was recorded and used to calculate stress and strain. A stress-strain curve was then generated using the following equations:

where *d* is diameter and *d_i_* is initial diameter.

$$\boldsymbol{Strain}\left( \boldsymbol{\lambda} \right)\boldsymbol{=}\frac{\boldsymbol{\Delta d}}{\left[ \boldsymbol{d}\left( \boldsymbol{i} \right) \right]}$$

where *L* is one-dimensional load, *H* is intima media thickness, and *D* is vessel length.

$$\boldsymbol{Stress}\left( \boldsymbol{t} \right)\boldsymbol{=}\frac{\boldsymbol{\lambda L}}{\left[ \boldsymbol{2}\left( \boldsymbol{HD} \right) \right]}$$

The elastic modulus of the stress-strain curve was determined as the slope of the linear regression fit to the final four points of the stress-strain curve, as previously reported by our laboratory^6^. Aortic intima media thickness and diameter were assessed as we have described previously^2^. Briefly, aortic rings (1mm) were frozen in optimal cutting temperature solution and stored at -80°C until the time of sectioning. Aortic sectioning was performed on a Cryostat (Leica Biosystems, Wetzlar, Germany) at 22°C, and 7 μM sections were visualized, and images were captured with a bright-field microscope. Aortic intima media thickness and diameter were calculated using ImageJ software.

**Aortic protein abundance**

***Immunoblotting.*** Protein abundance was measured in segments of thoracic aorta following mechanical homogenization in a bullet blender (Next Advance, Troy, NY) with zirconium oxide beads (3:1 1mm:0.5mm beads, Next Advance, Troy, NY) in a radioimmunoprecipitation assay lysis buffer supplemented with protease and phosphatase inhibitors [1 mMol/L sodium orthovanadate, 1X complete mini protease inhibitor cocktail tablet (Roche, Mannheim, Germany), 1 mMol/L phenylmethylsulfonyl fluoride, 1:100 Phosphatase Inhibitor Cocktail (Sigma-Aldrich, St. Louis, MO), 5 mMol/L sodium fluoride, and 5 mMol/L sodium pyrophosphate. Total protein content was quantified using a bicinchoninic acid assay (Thermo Fisher Scientific, Eugene, OR). Next, abundance of Glyoxalase-1 (Glo1) (antigoat; 1:50; R&D Systems, cat no. AF4959) were determined by loading 0.4 μg/μL of aortic protein per capillary in a 25-lane (capillary) automated Western blot quantitative analyzer (JESS, ProteinSimple, San Jose, CA), according to the manufacturer’s guidelines, as described previously^1,6^, following the validation of these antibodies in test aorta lysates. Anti-goat and anti-rabbit secondary antibodies were provided by the manufacturer and used according to the manufacturer’s guidelines. A grayscale analysis of the band intensities was then performed to quantify protein abundance using Compass software (ProteinSimple), with target proteins expressed relative to total protein.

***Immunofluorescence.*** Immunofluorescence assays were performed to visualize the subcellular localization of target proteins; Collagen-I and methylglyoxal derived hydroimidazolone-1 (MGH-1). At the time of sacrifice, ~1mm sections of thoracic aorta were excised and frozen in OCT (Tissue-Tek^®^ O.C.T.) compound, as described above^1,9^. Later in time, 7 μm sections (Leica CM1520) were plated on poly-L-lysine-coated microscope slides, fixed in 2-4% paraformaldehyde, washed with PBS, and permeabilized (0.1% Triton X-100). Slides were then incubated with anti-collagen-1 primary antibody (1:200; Southern Biotech, Birmingham, AL; Cat# 1310-01) for 1 hour, washed with PBS, and incubated with a species-specific fluorescent secondary antibody (AlexaFluor 647; Invitrogen, Waltham, MA) for 60 minutes. Slides were washed, stained with DAPI (1:1000; Invitrogen, Waltham, MA; Cat# D1306) for 5 minutes, and cured overnight with ProLong Gold mounting media (Invitrogen, Waltham, MA; Cat# P36980). For MGH-1, slides were incubated with anti-MGH-1 primary antibody (1:200; Cell Biolabs, San Diego, CA; Cat# STA-011) according to instructions from the mouse-on-mouse immunodetection kit (Vector laboratories, Newark, CA; Cat# BMK-2202). The slides were then imaged using EVOS m7000 (ThermoFisher, Waltham, MA; Cat# AMF7000) fluorescence microscope under identical conditions and analyzed using Invitrogen Celleste 5.0 Image Analysis Software. Images for were obtained at 10x magnification. The abundance of collagen-1, and MGH-1 was determined as the average intensity (A.U.) of the positively stained area across N = 6-8 samples/animal. Images used to evaluate elastin breaks were captured at wavelengths between 470 and 525 nm, utilizing the autofluorescence of elastic lamellae, as described previously^10,11^. Elastin breaks were identified as regions with substantial discontinuities in the elastic lamellae. To minimize errors caused by lamellae being out of the microscope's focal plane, breaks were confirmed across two to three images per animal. The number of elastin breaks was averaged from n = 2 or 3 samples per animal.

***Quantification of free MGH-1***

10 µL of serum was added to 200 µL of 80:20 MeOH:ddH_2_O (−80°C) containing ten pmol ^13^C-MG-H1 and extracted at −80°C overnight. Insoluble protein was removed via centrifugation at 14,000 x g for 10 min at 4°C, and supernatants were transferred to a new tube. 15 µL of heptafluorobutyric acid (1:1 in H_2_O) was added to each sample, and debris was removed via centrifugation at 14,000 × *g* for 10 min. Samples were analyzed as described previously^12^. (QuARKMod).

**Statistical Analysis**

Statistical analyses were conducted in Prism, version 10 (GraphPad Software, Inc. La Jolla, CA, USA). Data were first assessed for outliers (ROUT method, *Q* = 1%) and normality (Shapiro-Wilk normality test, *P* > 0.05) within groups. Two-way mixed ANOVAs were used to determine differences in aortic PWV, systolic and diastolic blood pressures (group x time [pre/post treatment]). Differences across animal groups in artery characteristics, immunofluorescence, and qPCR were assessed using one-way ANOVA. Plasma exposure-based changes in elastic modulus, immunofluorescence markers were assessed using paired samples t-tests as different media conditions were tested on arteries obtained from the same mouse. When significant main effects were detected, pairwise comparisons were made using the Holm-Sidak post hoc test. Significance was set to α = 0.05. Unless otherwise noted, data are presented as mean ± SEM.

12.	Wimer L, Kaneshiro KR, Ramirez J, Bose N, Valdearcos M, Shanmugam MM, Farrera DO, Singh P, Beck J, Sellegounder D, Enqriquez Najera L, Melov S, Ellerby L, Cho S-J, Newman JC, Koliwad S, Galligan J, Kapahi P. Glycation-lowering compounds inhibit ghrelin signaling to reduce food intake, lower insulin resistance, and extend lifespan

**Supplemental Table 1**. Characteristics of mice administered with doxorubicin (DOXO) or sham followed by treatment with the ABT-263 or vehicle

|  | Sham  Veh | Sham ABT-263 | DOXO  Veh | DOXO  ABT-263 |
| --- | --- | --- | --- | --- |
| *n* | 12  *3 female*  *9 male* | 11  *6 female 5 male* | 10  *4 female*  *6 male* | 11  *5 female*  *6 male* |
| Aorta |  |  |  |  |
| Diameter, μM | 616 ± 19 | 640 ± 20 | 631 ± 7 | 617 ± 18 |
| Intima media thickness, μM | 40 ± 1 | 39 ± 1 | 40 ± 2 | 42 ± 2 |
| Systolic blood pressure, mm/Hg |  |  |  |  |
| Pre | 99 ± 2 | 98 ± 3 | 101 ± 2 | 101 ± 3 |
| Post | 96 ± 3 | 95 ± 5 | 96 ± 2 | 95 ± 3 |
| Diastolic blood pressure, mm/Hg |  |  |  |  |
| Pre | 71 ± 2 | 71 ± 3 | 73 ± 2 | 76 ± 2 |
| Post | 69 ± 4 | 68 ± 4 | 69 ± 2 | 65 ± 3 |

Data are mean ± SEM.

**Supplemental Table 2.** Circulating concentrations of advanced glycation end products.

Data are Mean ± SEM. CEL (carboxyethyl-lysine); CEA (carboxyethyl-arganine); CML (carboxymethyl-lysine); MGO (methylglyoxal).

|  | **Sham-Veh** | **Doxo-Veh** | **Doxo-ABT263** | ***P* value** |
| --- | --- | --- | --- | --- |
| ***n*** = | 6 | 7 | 6 |  |
| **CEL (μMol)** | 0.297 ± 0.024 | 0.330 ± 0.052 | 0.330 ± 0.052 | 0.854 |
| **CEA (μMol)** | 0.118 ± 0.007 | 0.107 ± 0.013 | 0.107 ± 0.009 | 0.673 |
| **CML (μMol)** | 0.385 ± 0.036 | 0.435 ± 0.055 | 0.348 ± 0.044 | 0.443 |
| **MGO (μMol)** | 2.250 ± 0.096 | 2.267 ± 0.098 | 2.333 ± 0.102 | 0.822 |

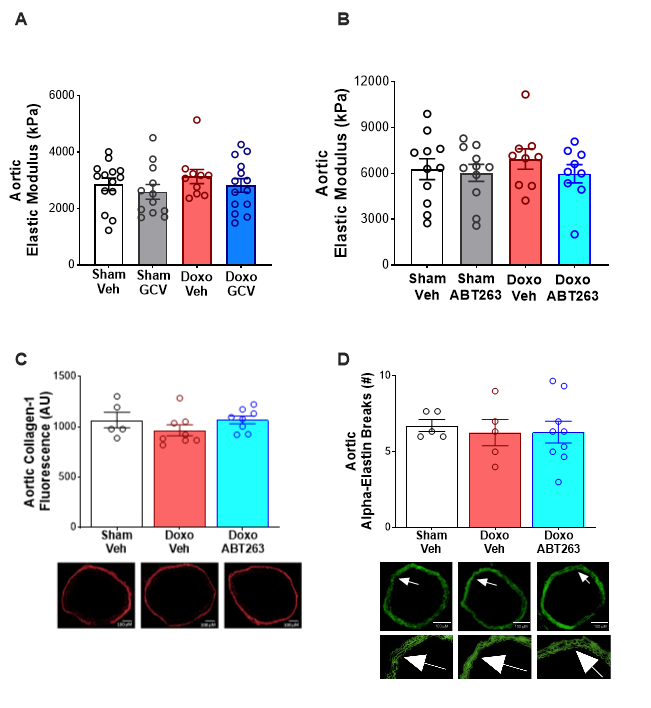

**B**

**A**

**Supplementary figure 1: Senolytic treatment after Doxo administration does not influence collagen deposition or elastin fragmentation**. Aortic collagen-1 fluorescence **(A)**. Aortic elastin breaks **(B)**. All values are mean ± SEM. n=5-9/group for (C) and (D).

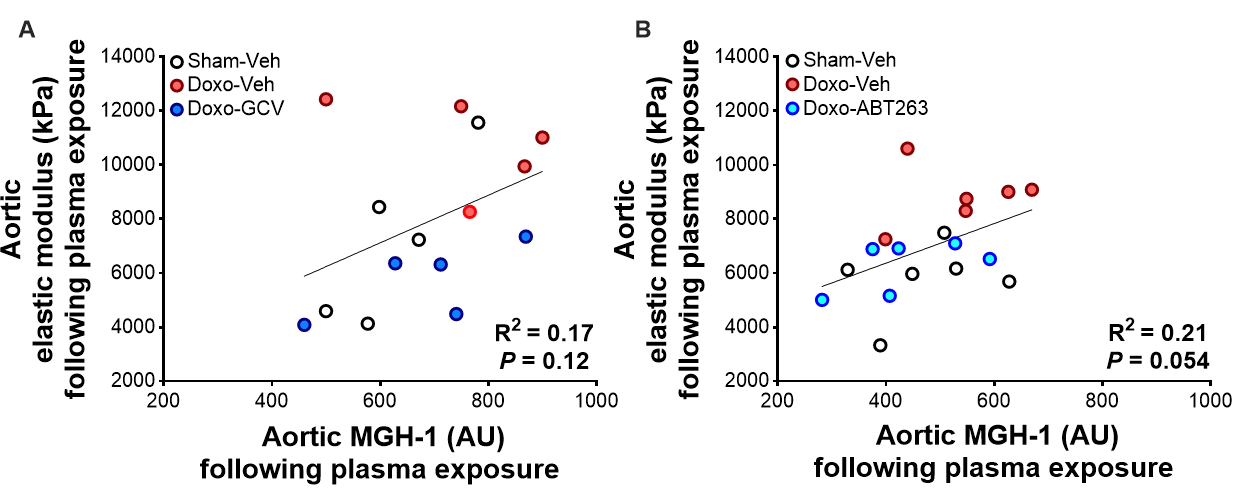

**Supplemtary figure 2: Aortic MGH-1 levels following plasma exposure are correlated with aortic elastic modulus**. Aortic MGH-1 levels relative to the aortic elastic modulus following plasma exposure from study 1 **(A)** and 2 **(B)**.

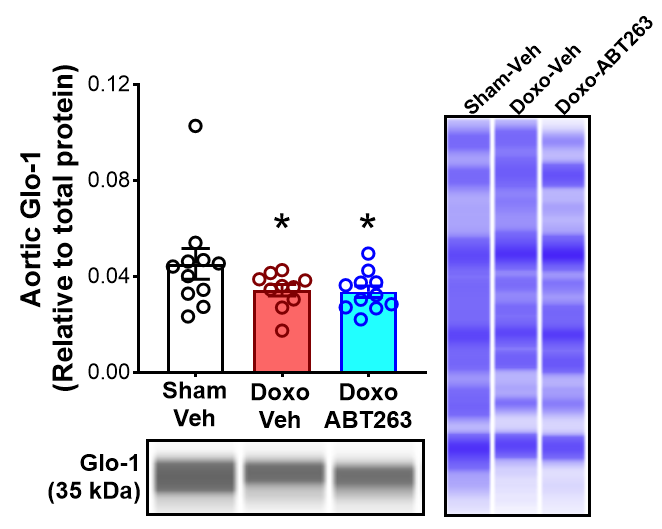

**Supplemtary figure 3: Doxo adminsitration lowers Glyoxalse-1 (Glo-1) levels and senolytic treatment has no effect.** Aortic glyoxalase-1 levels. All values are mean ± SEM. n=8-11/group.
